# Kidney mRNA-protein expression correlation: What can we learn from the Human Protein Atlas?

**DOI:** 10.1101/2024.07.12.601822

**Authors:** Dianne Acoba, Anna Reznichenko

## Abstract

The Human Protein Atlas (HPA), with more than 10 million immunohistochemical images showing tissue- and cell-specific protein expression levels and subcellular localization information, is widely used in kidney research. The HPA contains comprehensive data on multi-tissue transcript and protein abundance, allowing for comparisons across tissues. However, while visual and intuitive to interpret, immunohistochemistry is limited by its semi-quantitative nature. This can lead to mismatches in protein expression measurements across different platforms. We performed a comparison of the HPA’s kidney-specific RNA sequencing and immunohistochemistry data to determine if the mRNA and protein abundance levels are concordant. Our study demonstrates that there is a discordance between mRNA and protein expression in the kidney based on HPA data. Using an external validatory mass spectrometry dataset, we show that more than 500 proteins undetected by immunohistochemistry are robustly measured by mass spectrometry. The HPA transcriptome data, on the other hand, exhibit similar transcript detection levels as other kidney RNA-seq datasets. Such discordance in mRNA-protein expression could be due to both biological and technical reasons, such as transcriptional dynamics, translation rates, protein half-lives, and measurement errors. This is further complicated by the heterogeneity of the kidney tissue itself, which can increase the discordance if the cell populations or tissue compartment samples do not match. As such, shedding light on the mRNA-protein relationship of the kidney-specific HPA data can provide more context to our scientific inferences when we discuss renal gene and protein quantification.

## INTRODUCTION

The complexity of chronic kidney disease (CKD) necessitates a nonreductionist systems biology approach to understand its multifactorial biology, making high-throughput or omics studies an important pillar in kidney research.^1^ Reference datasets are essential for benchmarking healthy expression levels and the Human Protein Atlas (HPA) is invaluable as it provides both transcript and protein abundance information.

The HPA is immensely used in kidney research, with more than 2 000 CKD research articles citing HPA’s seminal publication in 2015.^2^ Common use cases of HPA include checking tissue- and cell-specific mRNA and protein expression levels, subcellular localization, and protein structure and interactions. Other existing resources also provide kidney transcript and protein expression data, however, the HPA is currently the only platform that contains multi-tissue transcript and protein abundance information, which uniquely allows for a head-to-head comparison between mRNA and protein levels. It is also continuously being updated with new information as technologies emerge.

The HPA’s renal transcriptome data is derived from deep sequencing of mRNA (RNA-seq) of frozen kidneys of nine individuals, six females (48 - 67 years old) and three males (46 - 78 years of age). The kidney samples are composed of heterogeneous cell populations, accounting for 60-80% tubules, 5-25% glomeruli, 5-10% fibroblasts and 5-25% other cell types.^2^ RNA-seq allows for global and unbiased transcript identification and quantification in a single high-throughput sequencing assay, compared to the limited and targeted nature of microarray technologies. With RNA-seq, isoforms and differential exon usage can be measured, sequencing depth and precision are better compared to earlier technologies, and absolute quantification is possible.^3^ On the other hand, the kidney protein expression dataset is from antibody-based protein profiling through conventional brightfield immunohistochemistry (IHC) for normal kidney tissue. Tissue microarrays are stained with DAB (3,3’-diaminobenzidine)-labeled antibodies and counterstained using hematoxylin. The kidney proteome is represented by samples from three individuals (not uniform across all proteins) and IHC provides semi-quantitative and relative measurements that include information on the staining intensity, quantity, and location.

Previous studies exploring mRNA-protein expression relationships in different human tissues, including the kidney, showed a surprising lack of strong correlations.^4–6^ Variability in transcript-protein correlation has been observed across tissues (within-gene correlation), and even across genes (across-gene correlation). The variability is postulated to be due to spatial and temporal mRNA dynamics, post-transcriptional and post-translational regulation, translation rates, protein synthesis constraints and delay, protein transport and half-lives, and potential measurement error and technical variability.^3,7^ The kidney is not exempt from this correlation discordance, primarily driven by the tissue’s complexity and highly heterogeneous cell populations.^8^ Tissue complexity leads to incorrect estimates of mRNA-protein correlations, if the fraction of different cell types in the samples are unmatched.^3^

In this study, we compare HPA kidney mRNA and protein expression data, which has not been done previously and utilizes a semi-quantitative proteomics dataset. Analyzing the kidney mRNA-protein relationship using HPA data will permit us to infer better biological insights and to interpret our research results better.

## METHODS

Kidney protein and transcript expression data were downloaded from the Human Protein Atlas v23.0 on February 1, 2024.^2^ The protein expression data provided antibody reliability (“Approved”, “Enhanced”, “Supported”, and “Uncertain”), semi-quantitative expression level (“High”, “Medium”, “Low”, and “Not detected”), and cell type (“bowman’s capsule”, “cells in glomeruli”, “cells in tubules”, “collecting ducts”, “distal tubules”, “proximal tubules (cell body)”, and “proximal tubules (microvilli)”) information for 13 467 proteins. Protein data was then filtered to exclude entries with “Uncertain” antibody reliability, and to only include genes with glomerular (“cells in glomeruli”) and tubular (“cells in tubules”) expression data, as the other cell types have limited protein measurements.

To classify transcript expression data from RNA-seq according to levels similar to the protein expression dataset, nTPM (normalized transcripts-per-million) values equal to zero were marked as “Not detected” and the remaining were divided into tertiles and consequently assigned as low, medium, or high. Transcript and protein data were plotted for comparison; genes and proteins with concordant and discordant mRNA-protein levels were identified.

As external reference datasets, kidney compartment-specific mass-spectrometry proteomics data from KPMP PXD033207^9^ and healthy control RNA-seq transcriptomics data from Levin et al (2020)^10^ were used. All analyses were performed in R 4.3.1.^11^

## RESULTS

In the transcript expression data, 20 162 transcripts are measured in the kidney. Out of these, 3 764 are considered not detected (nTPM = 0) and the high, medium and low expression tertiles have 5 481, 5 452, and 5 465 transcripts, respectively. High expression is defined as 13.3 to 52 063.1 nTPM, medium expression as 3.0 to 13.3 nTPM and low expression as 0.1 to 3.0 nTPM. A total of 11 157 mRNA-protein pairs in the glomeruli and 10 108 in the tubules are used in subsequent analyses. In the glomerular IHC data, 5 988 proteins are not detected, and 2 078, 2 240 and 851 proteins have low, medium, and high expression, respectively. For the tubular IHC data, 3 080 proteins have no expression detected, while 1 201, 3 666 and 2 161 proteins have low, medium, and high expression, respectively.

To visualize how the transcript and protein expression reported in HPA correlate, kidney mRNA expression data was plotted against both glomerular and tubular IHC data (**Figure 1A-B**). Approximately 24% (2 682 pairs) of the glomerular and 38% (3 808 pairs) of the tubular mRNA-protein pairs show expression abundance agreement, while there are 8 472 pairs (75.93%) with discordant protein-mRNA abundance in the glomeruli and 6 298 (62.31%) in the tubules (**Table 1**). Despite this, statistical testing via chi square test shows that mRNA and IHC data are not independent of each other, showing that the two are associated. Of particular interest to us are those with undetected protein levels but measurable mRNA expression and vice versa, those with unmeasured mRNA but detectable protein levels. In both the glomeruli and tubules, there are more undetected proteins but with observable mRNA expression. In the glomeruli, 4 953 proteins (44.39%) are undetected but have measurable mRNA and 2 122 in the tubules (20.99%). The number of proteins with IHC signals but no detectable mRNA expression are 76 (0.68%) in the glomeruli and 141 (1.39%) in the tubules.

**Figure 1.**
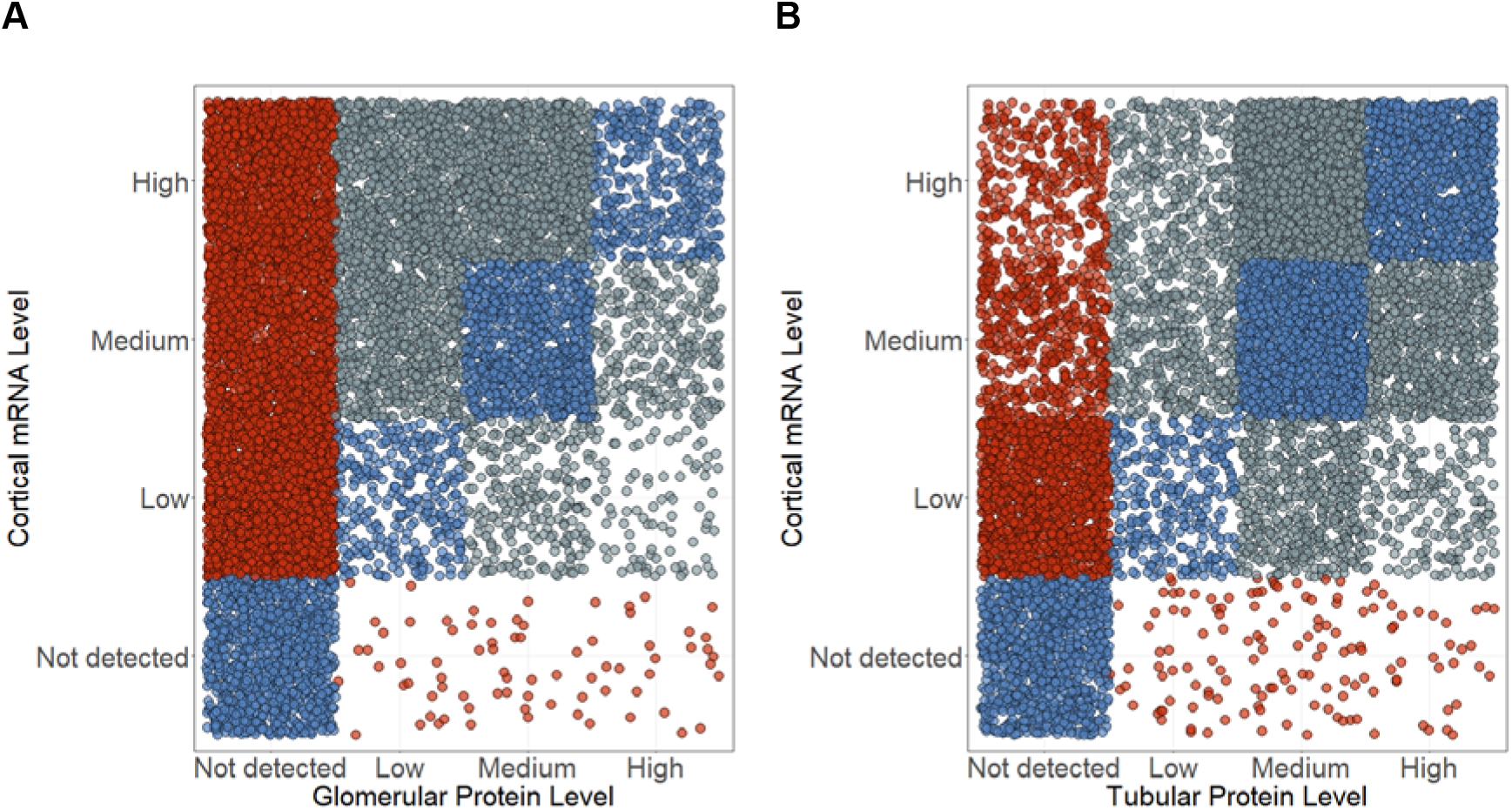
Kidney tissue mRNA & Protein abundances. Stripcharts demonstrate the correspondence between the protein and mRNA semi-quantitative expression levels in the (A) glomeruli and (B) tubules. Each individual dot represents one gene with both RNA and protein expression quantified. Perfectly concordant mRNA-protein pairs are shown with blue fill, the extremely discordant pairs are in red, and grey fill reflects the in-between scenarios.

**Table 1.** Count summary of the kidney proteins with discordant protein-mRNA abundance levels in the Human Protein Atlas.

|  | **Glomeruli** | **Tubulointerstitium** |
| --- | --- | --- |
| Number of reliable paired mRNA-protein expression measurements | 11 157 | 10 108 |
| mRNA-protein pairs with concordant abundance levels | 2 682<br>(24.04%) | 3 808 (37.67%) |
| mRNA-protein pairs with discordant abundance levels | 8 475<br>(75.96%) | 6 300 (62.33%) |
| Protein-mRNA pairs with no IHC signals but with detected kidney mRNA expression (nTPM > 0) | 4 953 | 2 122 |
| Protein-mRNA pairs with no IHC signals but with detected kidney mRNA expression (nTPM ≥ 1) | 3 773 | 1 172 |
| Protein-mRNA pairs with no IHC signals but with kidney mRNA expression detected (nTPM > 0) detected using compartment-specific mass-spectrometry data (mean spectral count ≥ 1) | 637 | 171 |
| Protein-mRNA pairs with no detectable mRNA but with IHC signals | 76 | 141 |
| Protein-mRNA pairs with no detectable mRNA but with IHC signals detected by compartment-specific RNA-seq (TPM > 0) | 62 | 120 |
| Protein-mRNA pairs with no detectable mRNA but with IHC signals detected by compartment-specific RNA-seq (TPM ≥ 1) | 5 | 3 |

Examples of extremely discordant pairs with respect to mRNA and protein expression are shown in **Figure 2**. *COL6A1*, which encodes for collagen type VI alpha 1 chain and is involved in extracellular matrix organization, is not detected in the glomeruli at the protein level through immunohistochemistry (**Figure 2A**) but has high transcript expression in the kidney (24.6 nTPM) and specifically, the glomeruli (18 TPM), according to HPA and Levin et al (2020), respectively. The same mRNA-protein abundance measure is observed for *HSP90AA1* encoding for the molecular chaperone heat shock protein 90 alpha family class A member 1 (**Figure 2B**) in the tubules (436.6 nTPM in HPA, 202 TPM in Levin et al). On the other hand, *H4C1*, encoding for H4 clustered histone 1, exhibits high protein expression as measured by IHC in both glomeruli and tubules (**Figure 2C**) but has very low to almost no detectable mRNA expression in both HPA and Levin et al (2020) RNA-seq data.

**Figure 2.**
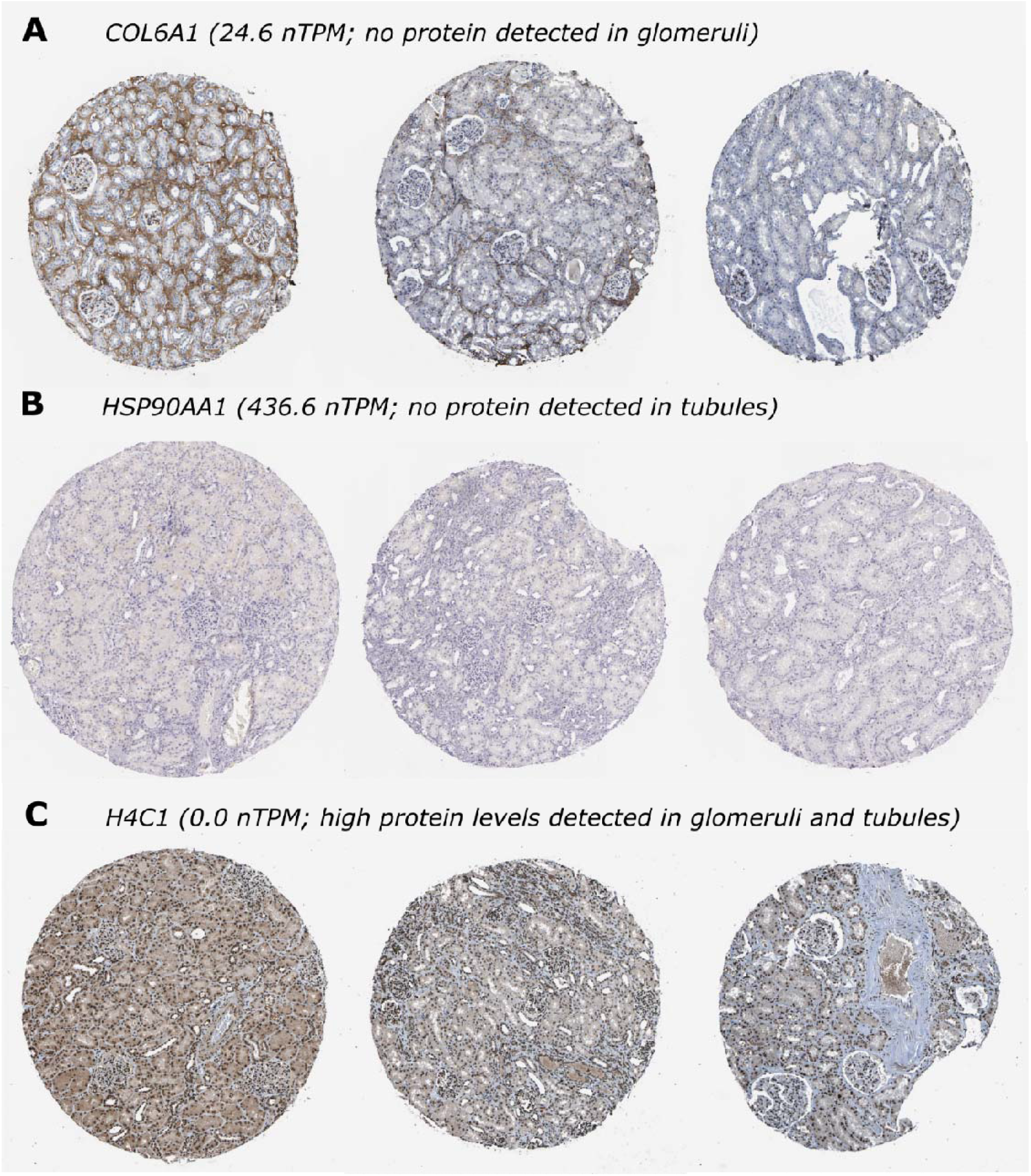
Representative immunohistochemical images of proteins with extremely discordant mRNA-protein expression levels according to the Human Protein Atlas. (A and B) Immunohistochemical images showing high glomerular COL6A1 (A) and tubular HSP90AA1 (B) protein expression. (C) Immunohistochemical images showing no detectable glomerular and tubular protein expression for H4C1. Image credit from the Human Protein Atlas. Images are available from https://v23.proteinatlas.org.

As a cross-platform validation analysis, we tested if the proteins with no IHC signals despite having mRNA expressed can be detected by mass spectrometry (MS). Using healthy glomerular and tubulointerstitial proteomics data, we determined that 637 of these IHC-undetected glomerular proteins and 171 tubulointerstitial proteins are detected by mass spectrometry with mean spectral count ≥1 (**Table 1**; **Supplementary File 1**). COL6A1 and HSP90AA1, despite having no discernible immunohistochemical staining (**Figure 2A-B**), have robust protein expression in the glomeruli (82 mean spectral count) and tubulointerstitium (97 mean spectral count), respectively, according to the KPMP MS data.

Out of the 76 glomerular and 141 tubular proteins with no detectable transcript expression, 62 and 120 have measurable mRNA expression based on Levin et al (2020) RNA-seq data, respectively. However, transcript expression is generally low, only 5 glomerular and 3 tubulointerstitial transcripts have mean TPM ≥ 1 (**Table 1**; **Supplementary File 1**).

## DISCUSSION

Immunohistochemistry allows for intuitive visual identification and localization of a target protein in both cellular and subcellular levels. It is routinely used in clinical practice for diagnostic pathology and is an important tool not just in healthcare but also in research. The HPA project has performed high-throughput IHC to map the human proteome in tissues and cells.^2^ However, the semi-quantitative nature and narrow range of staining intensity of IHC pose limitations in its usage. We hence set out to investigate if there is a disparity in the immunohistochemical and RNA-seq datasets in the HPA kidney-specific proteome and compare IHC-reported protein expression levels with kidney proteomics data.

We observed better concordance between the kidney mRNA and tubular IHC data than its glomerular counterpart, which could be due to the RNA-seq samples being 60 to 80% tubules. Kidney biopsies are mostly composed of cortical tissues, which primarily contain tubular cell populations. Cross-checking HPA’s IHC data with external mass spectrometry data, we found that some undetected proteins by IHC are expressed and detected in other datasets. On the other hand, HPA’s RNA-seq data seems to be more accurate as most of the undetected transcripts have also very low transcript expression in a validatory dataset. This is expected as the HPA RNA-seq data is mostly consistent with other human transcriptome datasets, such as GTEx and FANTOM5.^12^

Of interest are the mRNA-protein pairs with extremely discordant expression. In the first scenario where no mRNA is detected but protein expression is high, the discordance could be due to non-specific staining brought about by a promiscuous polyclonal antibody or IHC protocol failure, especially in the blocking step. The antibody could have low binding affinities leading to dissociation during processing steps.^13^ The epitope could also be located in a cellular compartment inaccessible to reagents or tissue artifacts could be present leading to false-positive staining due to leakage of proteins.^13^ In the tubular cells of the kidney, protein reabsorption happens and as such, could bind to antibodies non-specifically. For example, *H4C1* is highly expressed at the protein level according to IHC despite practically being undetected transcript-wise (**Figure 2C**) and HPA explains that this could be due to the antibody targeting proteins from more than one gene. The other scenario is when transcript expression is detected but immunohistochemistry does not detect any protein expression.

This could be due to a weak or diluted antibody and a failure in the antigen retrieval protocol failure.^13^ It could also be due to a difference in the expected location of the transcript and the protein, which is what is inferred about *COL6A1* (**Figure 2A**). These are just examples of how both biological and technical reasons could bring about such differences in abundance levels of the transcript and protein.

In this study, we only focused on the glomerular and tubulointerstitial data and excluded the others, which are a minority. However, this makes mRNA-IHC comparison even trickier due to potential variability in patient tissue samples and sample preparation differences. This is a limitation of both this study and HPA: the samples used for IHC and RNA-seq are different, thus, preventing us from comparing the same biosamples. Despite being considered histologically normal, the tissue samples are not from healthy individuals, who can have underlying disease that could affect kidney molecular processes and protein and transcript expression. Additionally, the HPA has a disclaimer on their website about how antibodies may not bind to its target due to differences in protein conformation and target accessibility. Protein denaturation and concentration, as well as sample complexity, are some of the factors that may influence off-target binding, which could lead to false results.^2^ HPA solves this issue by providing a reliability score to its data and also the reason why we chose to exclude all proteins with an “uncertain” reliability in this analysis.

Despite the depth of characterization housed by HPA, immunohistochemistry comes with limitations. Aside from its reliability being dependent on the antibodies used, it is also only a semi-quantitative measurement, manually interpreted by a pathologist, which can cause interobserver variability. Reproducibility also hinges on laboratory protocols. The most substantial hurdle is the narrow linear range of its staining intensity, resulting in rapid assay saturation.^14^ The single order of magnitude range of the antibody label DAB, for example, is poorly suited to assess markers across the full dynamic range of biological protein expression, which is approximately 8 orders of magnitude.^4^

Immunohistochemistry provides spatio-temporal expression information at a cellular or subcellular level, although at the price of quantitative measurements.^15^ Mass spectrometry methods can measure the dynamic protein expression range and is an important complement to IHC, by also providing isotype-specific information.^4,16^ However, MS has low sensitivity and is biased toward more precise detection of abundant proteins.^16^ The current development and improvement of high-throughput single-cell spatial proteomics methods, however, is one of the potential solutions to accurate protein quantitation.

We also want to discuss the possible biological reasons for discordant mRNA and protein levels aside from IHC technical limitations (e.g., antibody binding properties, interobserver variability in interpretation, narrow range of staining intensity). Other studies have discussed that discordant correlation between mRNA and protein levels can be due to regulatory elements that play diverse roles in translation.^4–6^ Previous studies have also reported that several mRNA elements affect translation and mRNA stability, such as codon usage, start codon context, and among others, secondary structures.^4^ Transcript range is also in four orders of magnitude, compared to eight in proteins, which explains the higher coverage of RNA-seq to MS. In addition, the number of protein molecules produced per mRNA molecule is much higher for abundant transcripts. It is postulated that genes encoding for abundant proteins have higher mRNA levels and also encode regulatory elements that lead to high translation efficiency and protein stability. On the other hand, small proteins are difficult to measure and MS sample preparation may also affect protein concentration.^4^

The Human Protein Atlas, with more than 10 million high-resolution IHC images in its portal, is a powerful tool for scientists engaging in protein and transcript expression studies.^15^ However, immunohistochemistry, while powerful in its ability for *in situ* protein detection at the single cell level, has its limitations which can greatly affect biological interpretation of scientific results. Our study shows how there is a discordance between the mRNA and protein abundance in the kidney dataset of HPA. We then validate the HPA IHC data with external mass spectrometry-based proteomics datasets and demonstrate that more than 500 proteins undetected by IHC are measured by mass spectrometry. Thus, we recommend scientists to exercise caution in treating HPA’s IHC data as the ‘ground truth’ for protein expression as it could have negative repercussions in research result interpretation. We must be careful and specific when discussing gene and protein quantitation, as they are not interchangeable.

## Data availability statement

All data analyzed in this study are publicly available and cited within the manuscript.

## Authors’ contributions

Dianne Acoba: Data curation, Formal analysis, Investigation, Methodology, Resources, Software, Validation, Visualization, Writing – original draft, Writing – review & editing Anna Reznichenko: Conceptualization, Funding acquisition, Project administration, Resources, Supervision, Writing – review & editing

